# HALO: A software tool for real-time head alignment in the MR scanner

**DOI:** 10.1101/2022.03.23.485491

**Authors:** Zhiying Zhao, Gigi Galiana, Cristofer Zillo, Terry Camarro, Maolin Qiu, Xenophon Papademetris, Michelle Hampson

## Abstract

Magnetic resonance imaging (MRI) studies in human subjects often require multiple scanning sessions/visits. Changes in a subject’s head position across sessions result in different alignment between brain tissues and the magnetic field which leads to changes in magnetic susceptibility. These changes can have considerable impacts on acquired signals. Therefore, we developed the Head Alignment Optimization (HALO) tool. HALO provides real-time visual feedback of a subject’s current head position relative to the position in a previous session. We verified that HALO enabled subjects to actively align their head positions to the desired position of their initial sessions. Our pilot sample of healthy subjects were able to improve their head alignment significantly using HALO and achieved good alignment with their first session meeting stringent criteria similar to that used for within-run head motion (less than 2mm translation or 2 degrees rotation in any direction from the desired position). Utilization of HALO in longitudinal studies will reduce the noise across sessions related to changes in magnetic susceptibility. HALO has been made publicly available.

## Introduction

In-magnet movement of the scanned tissues has been known as one of the most significant confounders to the functional MRI (fMRI) signal (Friston et al., 1996). Typical fMRI studies in humans usually consist of multiple runs or even sessions during which involuntary movement from the subjects is inevitable. Controlling for the artifacts arising from the intra-session head movements has been a convention in the field and a large body of tools are currently available for minimizing these artifacts prospectively (Zaitsev et al., 2017) or retrospectively (Friston et al., 1995). It is important to recognize that in addition to the local field strength, the blood oxygenation level dependent (BOLD) signal depends on the alignment of the brain tissues to the B0 field (Sati et al., 2012). Thus, a change in tissue alignment between MRI sessions can lead to changes in magnetic susceptibility gradients at the microscopic level which affect the apparent T2* (Bender and Klose, 2010) and cannot be controlled by the approaches mentioned above.

Against this background, we developed HALO for providing real-time visual feedback of head position from a previous session to subjects. It allows subjects to align their heads across sessions. The use of HALO in longitudinal MR imaging studies can reduce noise across sessions, particularly in regions of high susceptibility artifact.

We had a specific practical motivation for the development of HALO. We do real-time fMRI neurofeedback studies where the target pattern is defined in one session and trained in subsequent sessions, and we are interested in training regions of relevance to mental health that have high susceptibility artifacts (such as ventromedial prefrontal cortex and the amygdala ((Gerin et al., 2016; Scheinost et al., 2013)). Unfortunately, susceptibility artifacts can displace signal in the target area in a manner that is dependent on the alignment of the head. If head position is not controlled, a target region in an area of high susceptibility artifact will not be capturing signal representing the same neural substrate across sessions. This will undermine the efficacy of the neurofeedback.

In addition to the specific value for neurofeedback studies, HALO has general utility for improving signal quality in longitudinal MRI studies. Any longitudinal imaging study can use it to reduce the noise across sessions, particularly in areas of high susceptibility. High susceptibility regions such as the orbitofrontal cortex and the amygdala are of great interest in psychiatry and child development, therefore optimizing longitudinal studies to minimize noise in these areas is of general interest.

Finally, we suggest an exciting possibility (yet to be demonstrated) that by controlling head position across sessions (and also adjusting for field changes as monitored by field map collection), we may remove a current limitation of functional MRI: the inability to examine changes in brain activity across sessions using BOLD imaging.

## Methods

### Software design

The idea of HALO is to continuously collect the spatial information of the brain via EPI (Echo Planar Imaging) scans and present the displacement between the current position relative to the reference position (collected from the previous session, ‘target position’) to the subject in the form of visual feedback updated in near real-time, with which the subject performs voluntary movement to minimize the displacement.

HALO is developed in MATLAB (v2019a; Natick, Massachusetts: The MathWorks Inc.) with the graphical user interface (GUI) created using the App Designer functionality (Figure 1). During real-time alignment mode/online mode, the processing of a new volume consists of four steps:

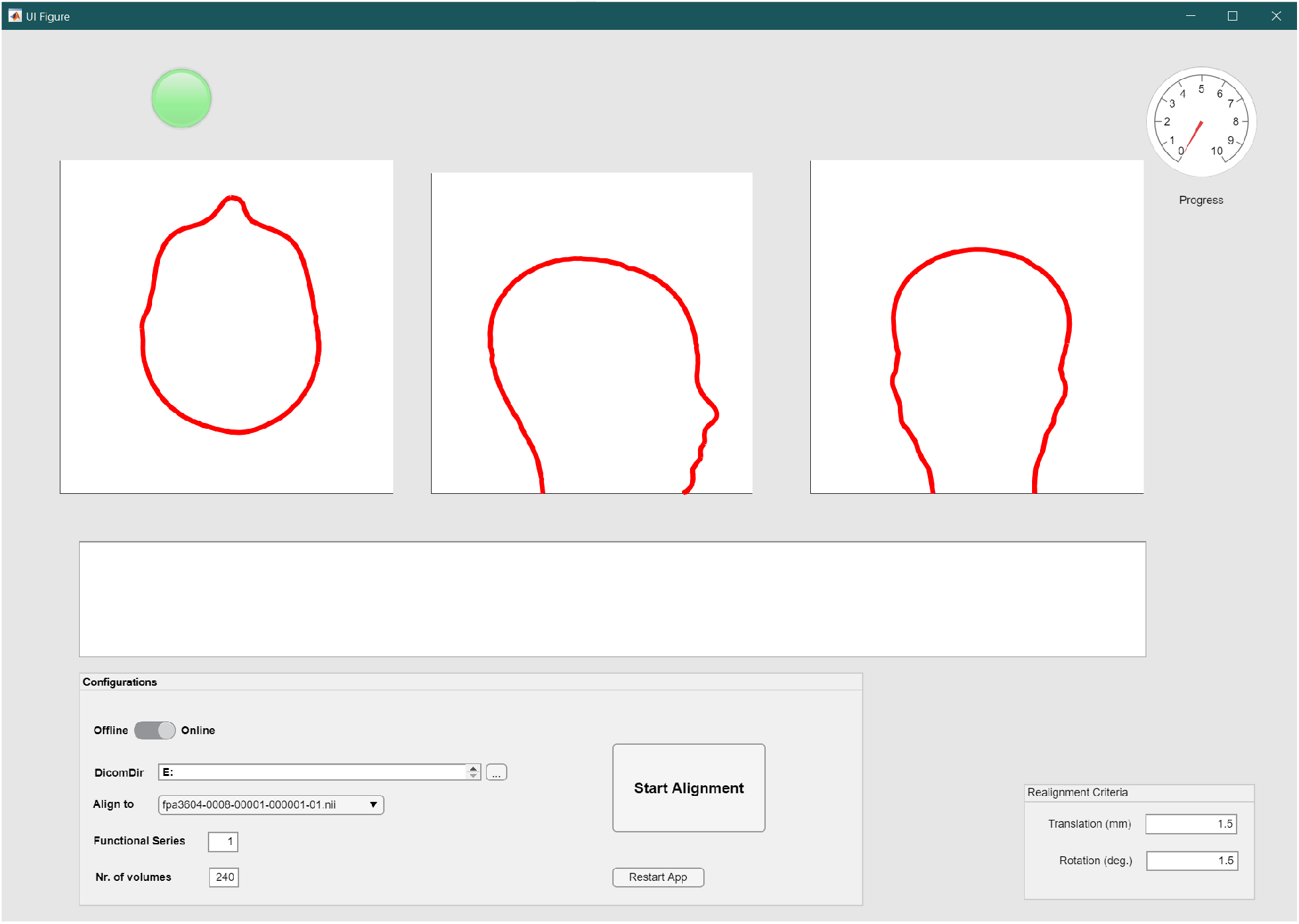

**Figure 2.**
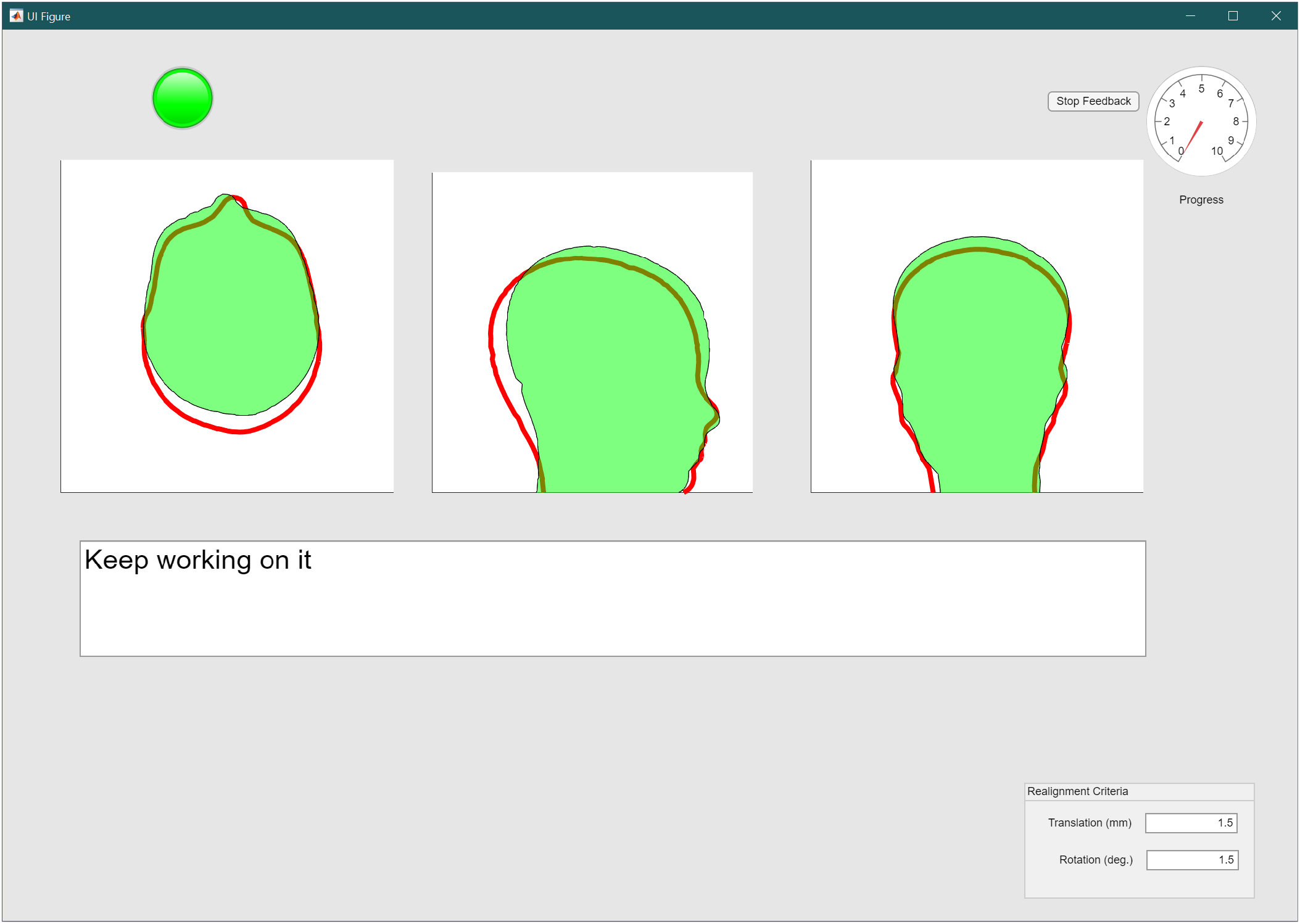
Interface of HALO during real-time alignment

1. Converting the DICOM image detected by the tool as a new volume to NIfTI format
2. Calculating the 4×4 rigid transformation matrix between the reference image and the new image which represents the rigid body movement from the target position to the current new position
3. Applying the transformation matrix to the virtual head object of the reference brain to allow visualization of the current head position relative to the reference brain
4. Visualizing both target and current position of the head on the interface and providing written as well as animation cues to indicate the progress of the alignment once the subject reaches the criteria set for sufficiently accurate alignment (see Figure 1 and Figure S1 in Supplementary Materials)

Scripts (namely *spm_dicom_convert*.*m* and *spm_realign*.*m*) from SPM12 (v7219; www.fil.ion.ucl.ac.uk/spm/software/spm12/) were separately adopted for step 1 and 3 for converting the Dicom images and calculating the 6-parameter rigid body movements (Friston et al., 1995) under the terms of the GNU General Public License. HALO is shared on https://osf.io/d4b59/.

### Tool test procedure

To evaluate the effectiveness of HALO in aligning head positions, 7 healthy subjects (4 males and 3 females) were recruited from the local community for the pilot study. None of the subjects had previous experience using HALO. Subjects were informed about the purpose and procedure of the study and written consent was obtained prior to participation. Each subject underwent two MRI sessions on a 3-Tesla Siemens Prisma system (software version syngo MR VE11C) using the Head/Neck 64 channel imaging coil. During the first session, an EPI volume was collected that they were instructed to align to during the alignment runs of the second session. The two imaging sessions were spaced one to two weeks apart for each subject. In both sessions subjects wore a pair of MRI-compatible headphones for communicating.

### Imaging Parameters

HALO was designed to update feedback every volume in the alignment runs. During tests before the formal data collection, it was noticed that EPI sequences with repetition time (TR) of 2000ms were too slow for the subject to track the head movement and hence made the alignment difficult. Based on the current efficiency of HALO in processing each volume (500-600ms for a cycle of all 4 steps), we used a multiband EPI sequence with the multiband acceleration factor set to 2. This allows for collecting 37 slices every 800ms with echo time = 30ms, acquisition matrix = 64*64, flip angle = 80°, voxel size = 3.12*3.12*4.6mm^3^.

### Scan setup

To ensure the comfort of the subjects, fMRI researchers usually place a pillow inside the head coil. However, the nature of the pillow is important in controlling variations in the head elevation across sessions. When scanning the third subject in this study, it was found that the pillow (nylon cover stuffed with compressible foam) had the filling more compressed under the participant’s head on the second session such that the subject needed to lift her head above the pillow to match the higher position from the first session. This could not be maintained for a long duration. To avoid this problem for the last four subjects we used a 12.7mm-thick polyurethane foam sheet as a new head support material. This thin material has less potential for long-term deformation while providing acceptable head support.

Because the SPM realignment script assumes images from different frames are acquired in the same Field of View (FoV), to assure the successful alignment in HALO translates to physical space, it’s critical to ensure the FoVs are consistent across sessions in terms of its isocenter, boundaries and tilting angle. This was achieved in two steps. First, the MRI isocenters were assigned to the same location across sessions by consistent table positioning using the display on the scanner which shows the table position in millimeters. Second, after collecting the target volume, the EPI sequence was saved into a new scan protocol (“Program” on Siemens systems) which retained the information for FoV placement from the first session. This Program was then reused in the second session to recover the FoV placement which took MRI isocenter as its origin. For confirmation that we were obtaining consistent FoV placement, we used three water capsules attached to the inner surface of lower part of the head coil and verified their consistent position in the images obtained (see, e.g., Supplementary Figure S2).

### Head alignment

Prior to the second sessions, the subjects were familiarized with the feedback interface and were shown examples to further their understanding about what head movements they needed to perform to align their head position.

During the alignment runs (240 EPI volumes each), they were instructed to start aligning their head (in green) to the target head (outlined in red which represented head position in the first session). The text box area underneath was also updated in real-time to indicate the current alignment progress. After the displacement parameters reached the preset criteria on the bottom right corner, verbal instructions were given in the text box to inform the subjects that they could start adjusting their body posture to make themselves comfortable while trying to maintain their head position (Figure S1) (this verbal instruction would be removed if the head was moved out of the position during the body adjustment process and they would have the chance to align their heads again with the more comfortable body posture). In the meantime, the needle on the gauge meter started rotating to the right by one step for each volume acquired and continued as long as the head position remained within criteria. The alignment was considered completed when the needle reached 10, and the scan was then terminated by the experimenter.

### Data analysis

During offline data analysis, the displacements between the aligned head and target head (6 parameters: translations and rotations in x, y, z axes) were calculated using the same realignment function as online (*spm_realign)*. For each subject, two sets of such parameters were separately calculated for the first volume collected during alignment and the last volume collected during alignment against the target position. For translations and rotations, the absolute values of all three axes were averaged separately for before and after alignment. These translation and rotation numbers were then compared between the two timepoints using paired t-tests to confirm the improvement in alignment.

## Results

6 out of the 7 subjects were successful in achieving and maintaining alignment (the subject who experienced the pillow problem could not maintain it). On average these six subjects spent 11.17 minutes (SD = 2.83min) to complete the task. This time was calculated from the total number of volumes of the alignment runs. Their improvements in translations and rotations are illustrated in Figure 3.

**Figure 3.**
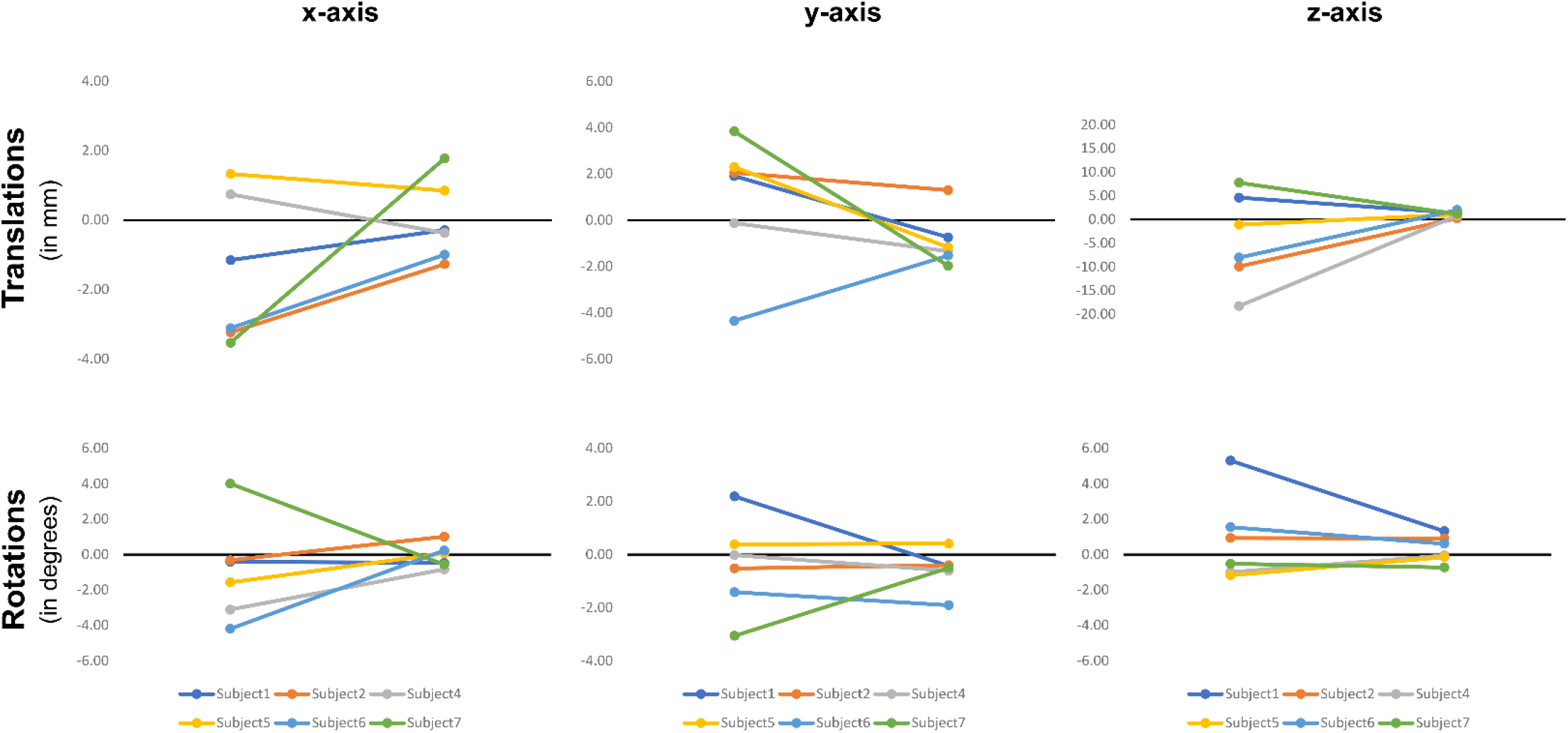
Change in translations along each axis (upper row) and rotations around that axis (lower row). Values before and after alignment are denoted by dots on the left and right respectively in each panel.

Results from the paired t-tests on the six subjects who completed the alignment task showed that translations were significantly reduced after alignment at the group level (t_5_ = 4.383, p = 0.007) as well as rotations (t_5_ = 3.473, p = 0.018). Results are still significant if we include changes in position for the third subject who was unable to maintain their alignment due to lack of head support (translation: t_6_ = 4.919, p = 0.003; rotation t_6_ = 3.119, p = 0.021). The full data for all subjects are provided in Supplementary Table S1. Individual data are shown for each movement dimension of each subject who completed in Figure 3.

To further demonstrate the efficacy of HALO in improving the alignment of anatomical structures, for each subject we also used the changes in centroid coordinates of left and right Brodmann Area 10 (BA10, the target brain structure in our OCD neurofeedback study (Scheinost et al., 2013)) after alignment as a real-world metric to measure the improvement in positioning. This metric is calculated by subtracting the BA10 centroid displacements (Before – After) using the centroid from Day1 as reference. The results indicate that after alignment, both left and right BA10 were closer to their positions in Day1 for every subject in our data (Table S2; see Supplementary Materials for the details of this calculation).

## Discussion

Aiming to solve the inter-session head alignment problem in the MRI environment, we developed HALO to help the subjects to align their heads voluntarily through movements guided by real-time feedback. In our pilot study, we showed that naïve subjects were able to reach the alignment criteria in less than 15 minutes with feedback from HALO.

HALO should be helpful for obtaining clean data in longitudinal imaging sessions. One specific longitudinal application we are interested in is multi-session real-time fMRI neurofeedback protocols that target brain regions with high susceptibility artifacts. However, any longitudinal imaging protocol may benefit from the use of HALO for minimizing changes in susceptibility across sessions.

Furthermore, the ability to keep susceptibility gradients consistent across imaging sessions may open new avenues for MRI research. For example, it may prove feasible to directly contrast brain activity across sessions using BOLD imaging if the local field inhomogeneities can be kept sufficiently stable by preserving head position. To be more specific, in conjunction with adjustments in the field homogeneity across sessions (based on field maps collected in each session), our alignment procedure may stabilize local field inhomogeneities sufficiently to enable meaningful contrasts in the BOLD signal across sessions. This would remove a basic limitation of BOLD imaging and open the door for a variety of psychiatric applications. For example, pharmacological studies examining the effects of a slow-acting drug on brain activity could use BOLD imaging not only to compare functional connectivity patterns on and off the drug, but to compare BOLD activity directly across sessions to examine if the baseline neural activity patterns in specific brain areas are altered by the drug.

A second novel application of HALO is in diagnostic structural imaging that uses image subtraction approaches (e.g., subtraction MRI). By matching the tissue-magnetic field alignment across sessions with the aid of HALO before structural scans, we hope to obtain cleaner subtracted structural images with which, for example, the development of brain lesions can be assessed more accurately.

A third potential application is for EEG-fMRI studies, because the magnetic induction induced artifacts have varied impacts on the EEG signals depending on the location of the electrodes in the magnet (Debener et al., 2008; Yan et al., 2009). HALO may help to eliminate the confounding effects from changes in electrode positions through alignment.

To be clear, more work is needed to determine the impact of head alignment across sessions in signal quality in the three novel applications described above. However, they represent exciting avenues for further development and are described here to illustrate the potential of HALO to have a substantial impact on the field of MR imaging.

Finally, there are several limitations of HALO. First, it requires real-time imaging capabilities. Second, although our subjects were able to align their head positions with the help of it, special measures were adopted to facilitate successful head alignment. These measures included controlling for head elevation change caused by pillow deformations and alignment of table and FoV positions across sessions. Furthermore, to allow voluntary head movement, unlike many other MRI studies, we did not insert cushions between the head and head coil. Therefore it remains to be evaluated how well the aligned head positions can be maintained in the subsequent imaging tasks without these cushions. Mitigating this concern somewhat was the presence of headphones on our subjects, that to some extent prevent involuntary movements. The use of a light piece of tape across the forehead, as recommended in a recent publication (Krause et al., 2019), may provide enough flexibility for head alignment while also facilitating stability during the subsequent scan. Finally, the time it took for head alignment may be an issue in imaging studies with limited time. We are confident that faster alignment times can be obtained, both by familiarizing participants with HALO (our study used only naive participants) and by optimizing the feedback time of it via further development. A variety of work needs to be done to develop the full potential of this line of research and HALO has been shared publicly to enable other researchers to both facilitate this development and to adopt it for use in their studies.

## Conclusion

We have developed a tool - HALO to enable subjects to align their head position in the magnet between sessions using real-time visual feedback. In our pilot study, naive subjects were able to improve their head alignment using it and to achieve a level of alignment across sessions similar to the alignment typically required within a single scanning run. Utilization of HALO may allow neuroimaging studies to assess longitudinal brain activity and structural changes more accurately and should improve multi-session fMRI neurofeedback studies that target brain areas with high susceptibility artifact. Future work is needed for further development of novel applications as discussed above.

## Acknowledgements

We thank Benjamin Thompson for naming HALO.

