## Supplemental Materials for "HALO: A software tool for real-time head alignment in the MR scanner"

Supplementary Materials

Figure S1. Interface when subject completes the alignment

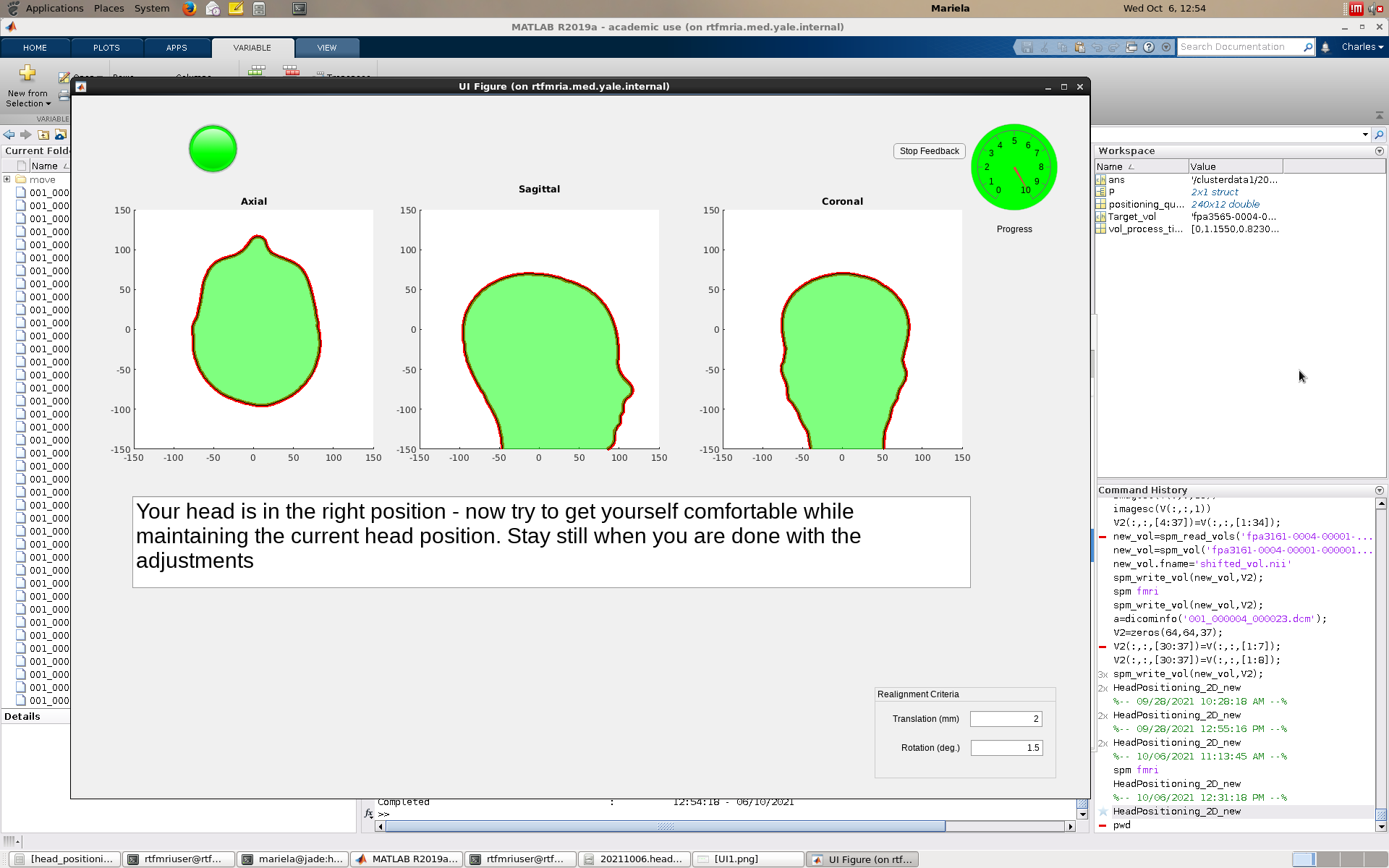

Figure S2. Axial view of brain images from one subject acquired with proton density sequence with the image taken after alignment (in red) overlaid on the image from the first session (in greyscale). The round objects on both side of the brain were two of the three water capsules (the third one only appears in higher axial slices) attached on the head coil from each session. These overlapped water capsules suggested that field of views remained stationary across sessions.

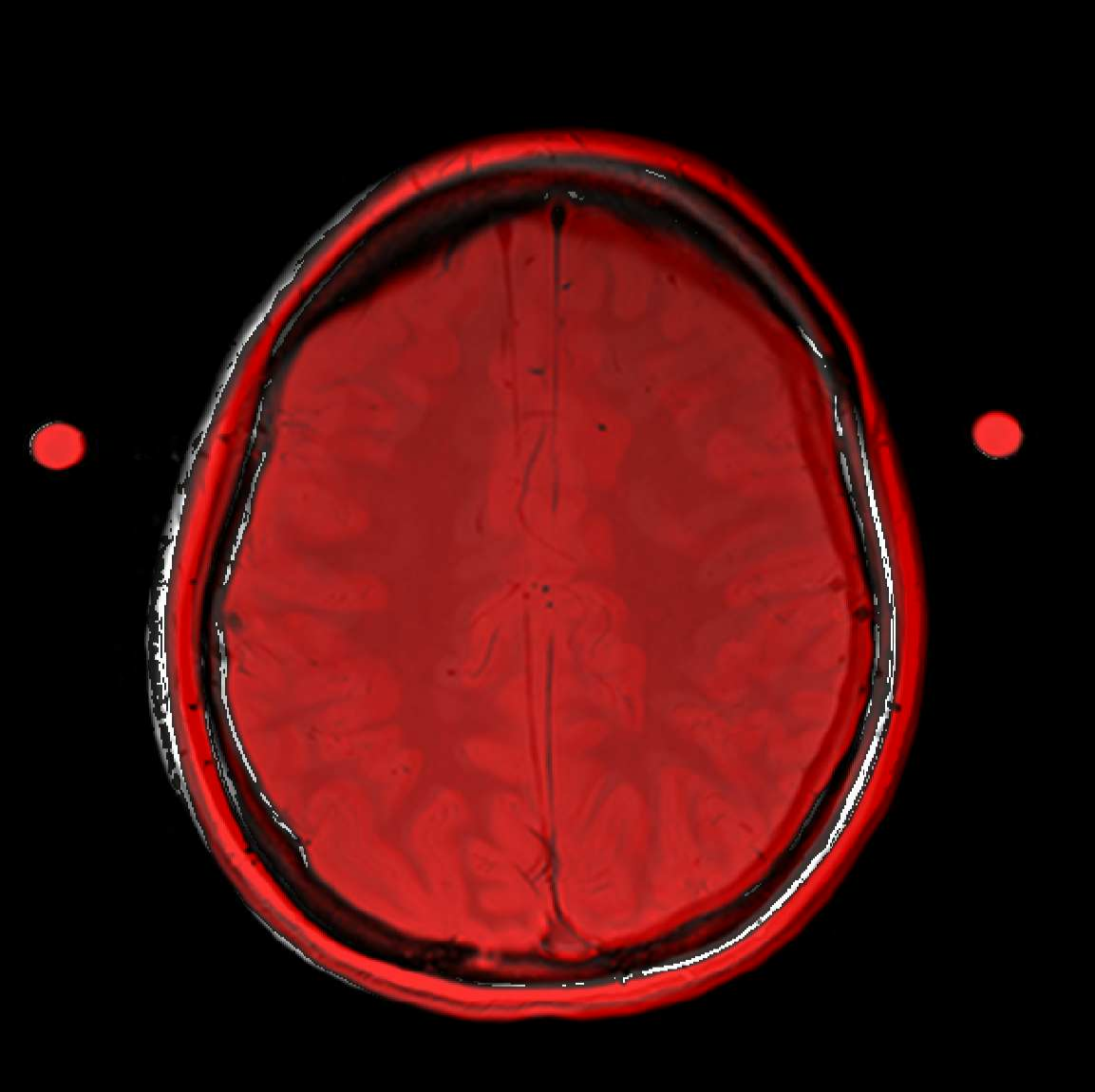

Table S1. Alignment results of all 7 subjects

|  | Translations in millimeters  (before alignment) | | | | Translations in millimeters  (after alignment) | | | | Rotations in degrees  (before alignment) | | | | Rotations in degrees  (after alignment) | | | |
| --- | --- | --- | --- | --- | --- | --- | --- | --- | --- | --- | --- | --- | --- | --- | --- | --- |
|  | x | y | z | Mean Abs. | x | y | z | Mean Abs. | x | y | z | Mean Abs. | x | y | z | Mean Abs. |
| Subject1 | -1.15 | 1.91 | 4.63 | **2.56** | -0.29 | -0.74 | 1.39 | **0.80** | -0.40 | 2.19 | 5.31 | **2.63** | -0.47 | -0.43 | 1.32 | **0.74** |
| Subject2 | -3.23 | 2.05 | -9.90 | **5.06** | -1.26 | 1.30 | 0.27 | **0.94** | -0.32 | -0.52 | 0.94 | **0.59** | 1.01 | -0.40 | 0.92 | **0.78** |
| Subject3 | 0.10 | -13.05 | -11.33 | **8.16** | 1.55 | 4.32 | -0.99 | **2.28** | -8.19 | 1.97 | 5.28 | **5.15** | 0.97 | 1.33 | -1.05 | **1.12** |
| Subject4 | 0.74 | -0.13 | -18.27 | **6.38** | -0.37 | -1.34 | 0.81 | **0.84** | -3.11 | -0.02 | -0.99 | **1.37** | -0.83 | -0.61 | -0.05 | **0.50** |
| Subject5 | 1.33 | 2.30 | -1.09 | **1.57** | 0.85 | -1.17 | 1.06 | **1.03** | -1.57 | 0.39 | -1.16 | **1.04** | 0.07 | 0.41 | -0.14 | **0.21** |
| Subject6 | -3.11 | -4.34 | -8.03 | **5.16** | -0.99 | -1.53 | 1.99 | **1.51** | -4.20 | -1.42 | 1.55 | **2.39** | 0.23 | -1.91 | 0.61 | **0.92** |
| Subject7 | -3.53 | 3.84 | 7.77 | **5.05** | 1.77 | -1.97 | 1.17 | **1.64** | 4.01 | -3.06 | -0.52 | **2.53** | -0.56 | -0.50 | -0.72 | **0.60** |

Table 1. The rigid-body movement parameters calculated for all the subjects using the target volume collected in the first sessions as the reference positions presented in the order of x-y-z axes. The averaged absolute translation/rotations before and after alignment are reported in the Mean Abs. columns and were used in the paired t-tests. Subject3 failed to reach alignment criteria because of the pillow problem.

Table S2. Improvement in BA10 centroid positions

|  | Left BA10 center change (in millimeters) | Right BA10 center change  (in millimeters) |
| --- | --- | --- |
| Subject1 | 1.19 | 2.33 |
| Subject2 | 8.58 | 8.90 |
| Subject3 | 15.76 | 9.44 |
| Subject4 | 15.41 | 14.05 |
| Subject5 | 1.59 | 1.95 |
| Subject6 | 5.72 | 1.53 |
| Subject7 | 8.61 | 1.91 |

**Calculation of the centroid changes**

The metrics in Table S2 was calculated in the order of registering the anatomical masks of bilateral BA10 to the individual functional space in Day1 (MNI→MPRAGE→EPI), extracting the Day1 centroid coordinates in world-space (in millimeters with the EPI FoV isocenter as the origin), calculating the new coordinates before and after alignment by multiplying the Day1 centroid coordinates by the rigid transformation matrices from Step 2, calculating the distances between the centroids (before alignment to Day1; after alignment to Day1) then subtracting the after alignment measure from the before alignment measure.
